# Vaccine-induced, but not natural immunity, against the Streptococcal Inhibitor of Complement protects against invasive disease

**DOI:** 10.1101/2020.11.22.393389

**Authors:** Lionel K. K. Tan, Mark Reglinski, Daryl Teo, Nada Reza, Lucy E. M. Lamb, Vaitehi Nageshwaran, Claire E. Turner, Mats Wikstrom, Inga-Maria Frick, Lars Bjorck, Shiranee Sriskandan

## Abstract

Highly pathogenic *emm1 Streptococcus pyogenes* strains secrete the multidomain Streptococcal inhibitor of complement (SIC) that binds and inactivates components of the innate immune response. We aimed to determine if naturally occurring or vaccine-induced antibodies to SIC are protective against invasive *S. pyogenes* infection. Immunisation with full length SIC protected mice against systemic bacterial dissemination following intranasal or intramuscular infection with *emm1 S. pyogenes.* Vaccine-induced rabbit anti-SIC antibodies, but not naturally occurring human anti-SIC antibodies, enhanced bacterial clearance in an ex vivo whole blood assay. SIC vaccination of both mice and rabbits resulted in antibody recognition of all domains of SIC, whereas naturally occurring human anti-SIC antibodies recognised the proline-rich region of SIC only. We therefore propose a model whereby natural infection with *S. pyogenes* generates non-protective antibodies against the proline-rich region of SIC, while vaccination with full length SIC permits development of protective antibodies against all SIC domains.

## Introduction

Invasive disease caused by the human specific pathogen, *Streptococcus pyogenes*, also known as group A Streptococcus (GAS), has been increasing since the 1980s and is associated with mortality of approximately 20%^1,2^. Strains expressing the M1 protein, encoded by *emm1,* are overrepresented amongst invasive isolates, and account for over 30% of cases of necrotising fasciitis and streptococcal toxic shock syndrome ^3^. The Streptococcal Inhibitor of Complement (SIC) is an extracellular protein, almost uniquely expressed by *emm1 S. pyogenes,* and is one of several virulence factors implicated in the propensity for *emm1* isolates to cause severe infection^4^.

Three distinct regions of SIC have been described; an N-proximal short repeat region, a central long repeat region, and a C-proximal proline-rich region ^4,5^. SIC binds to the C5b67 complex of complement to inhibit the formation of the membrane attack complex ^4,6^. SIC also inhibits the function of host antimicrobial factors including lysozyme, alpha and beta defensins, secretory leucocyte protease inhibitor, LL-37 and histones, and additionally has a role in bacterial adherence to epithelial cells ^5,7-10^. While transcriptomic and mutagenesis studies have suggested a role for SIC in invasive disease *in vivo*^11,12^, expression of SIC has not been directly linked to invasiveness of *S. pyogenes* in the clinical setting. Recently SIC was one of the 15 streptococcal proteins detected in pleural fluid from a child with empyema caused by *emm1 S. pyogenes,* indicating that SIC is expressed at high levels during natural infection although it may be degraded ^13^.

Despite a relatively low incidence of anti-M1 antibody, antibody to SIC is found frequently (~40%) among humans from diverse populations, including healthy people and also those with previous streptococcal disease ^14-16^. SIC is genetically highly variable with over 300 *sic* alleles described and it is speculated that human antibody at mucosal surfaces may drive SIC variation ^17^, however the role of anti-SIC antibodies in host immunity remains unclear. We set out to measure SIC production by *S. pyogenes in vitro* and *in vivo,* and then determine whether immunity to SIC can be protective. We found that, despite the prevalence of naturally occurring anti-SIC antibodies in humans, these antibodies do not confer opsonophagocytic protection against *S. pyogenes.* In contrast, vaccine-induced antibodies against full length SIC do confer opsonophagocytic activity against *S. pyogenes* and, furthermore, provide protection against experimental invasive streptococcal disease.

## Results

### Expression of SIC in vitro among invasive and non-invasive isolates

SIC expression in broth was quantified by western blot and densitometry from 101 clinical isolates of *emm1 S. pyogenes* to determine whether SIC expression was associated with site of bacterial isolate or original disease phenotype (Figure 1). SIC expression varied from 4.14 ng/ml to 434.67 ng/ml (median 83.68 ng/ml, IQR 45.43-126.63) in culture supernatant. Although there was a wide range of expression, there was no significant difference in the detected levels of SIC expression between invasive disease isolates (median 80.58 ng/ml, IQR 43.92-118.4), and non-invasive isolates (median 88.06 ng/ml, IQR 42.69-150.7) (Figure 1A). Further categorisation of the 87 strains for which the site of isolation was known did not reveal any association between SIC expression and any specific disease etiology (Figure 1B). Among a subset of 39 isolates for which the sequence of the negative regulatory locus covRS was known, SIC secretion *in vitro* was higher in the 6 strains with covRS mutations (median 311.8 ng/ml) than strains without mutations (median 88.06 ng/ml, p=0.0017).

**Figure 1.**
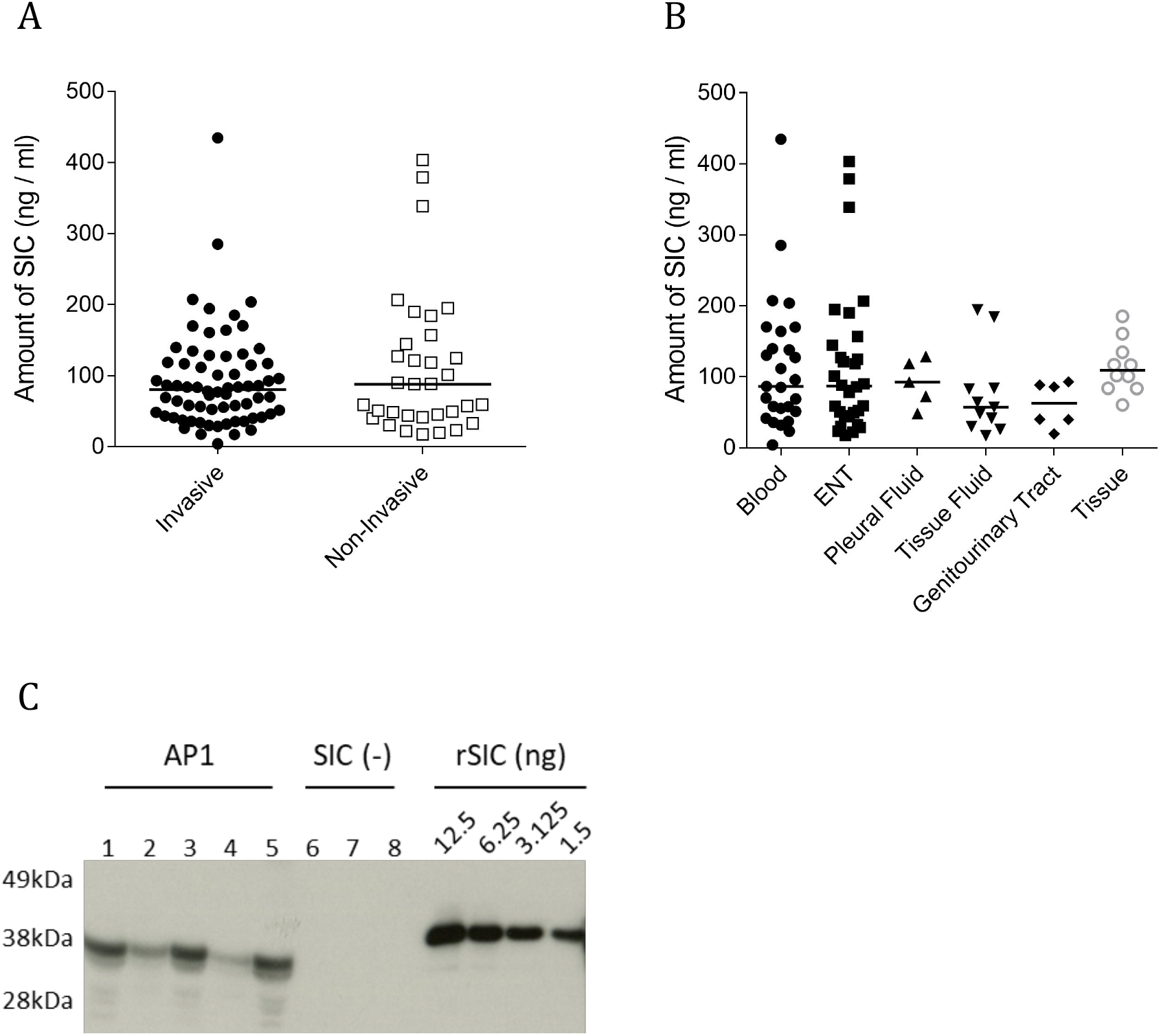
Quantification of *in vitro* and *in vivo* SIC production. A-B) The concentration of SIC in overnight culture supernatants from 101 *emm1 S. pyogenes* clinical isolates grouped by A) invasive vs non-invasive disease phenotype or B) site of isolation was quantified. (C) SIC was quantified in the thigh tissue of mice following a 3 h intramuscular infection with the *emm1* GAS isolate AP1 (5 mice, lanes 1-5) or a SIC-negative AP1 derivative (3 mice, lanes 6-8). Quantifications were performed by Western blotting and densitometry using a recombinant SIC (rSIC) standard ranging from 12.5 ng to 1.56 ng per well.

To quantify SIC expression *in vivo*, FVB/n mice were infected intramuscularly with 5 × 10^7^ CFU of *S. pyogenes emm1* strain AP1 or SIC-negative derivative of AP1 ^9^. SIC was detected in infected muscle tissue extracts from AP1-infected mice (n=5) three hours post-infection (Figure 1C); the median quantity of bacteria was 3.6 × 10^7^ CFU (0.9 to 6.5 ×10^7^ CFU) per mg of thigh tissue and the median SIC level detected was 2.49 ng/mg of tissue (0.39 to 3.27 ng/ml of tissue). SIC was not detected in thigh tissue of mice infected with the SIC-negative derivative (n=3); the median quantity of bacteria was 6.6 × 10^6^ CFU (6.5 × 10^6^ to 1.47 ×10^7^ CFU) of SIC-negative derivative per mg of thigh tissue (Figure 1C).

### SIC is immunogenic in mice and immunisation improves outcome following intranasal infection

Serum IgG antibodies directed against SIC are common in healthy human populations ^14-16^, but little is known about the protective role of these antibodies against disease. The potential protective role of anti-SIC antibodies was therefore assessed in a mouse model of infection. To determine whether recombinant SIC protein variant SIC1.300 (rSIC1.300) was immunogenic, mice were immunised with rSIC1.300 or sham vaccine containing PBS and adjuvant. Following immunisation, anti-SIC antibodies could be detected in a 1:64,000 dilution of immunised mouse sera (Figure 2A). To assess whether these titres translated into protective immunity, immunised mice were infected with a contemporary *emm1 S. pyogenes* strain H584 (a strain that expresses the SIC1.300 variant) via the intranasal route using a volume known to reach the lung and disseminate systemically. *S. pyogenes* lower respiratory tract infection led to noticeable systemic disease. However, mice immunised with recombinant SIC1.300 demonstrated improved outcomes (time to humane endpoints) compared to mice immunised with sham vaccine (Figure 2B), even when the challenge dose was increased (Figure 2C).

**Figure 2.**
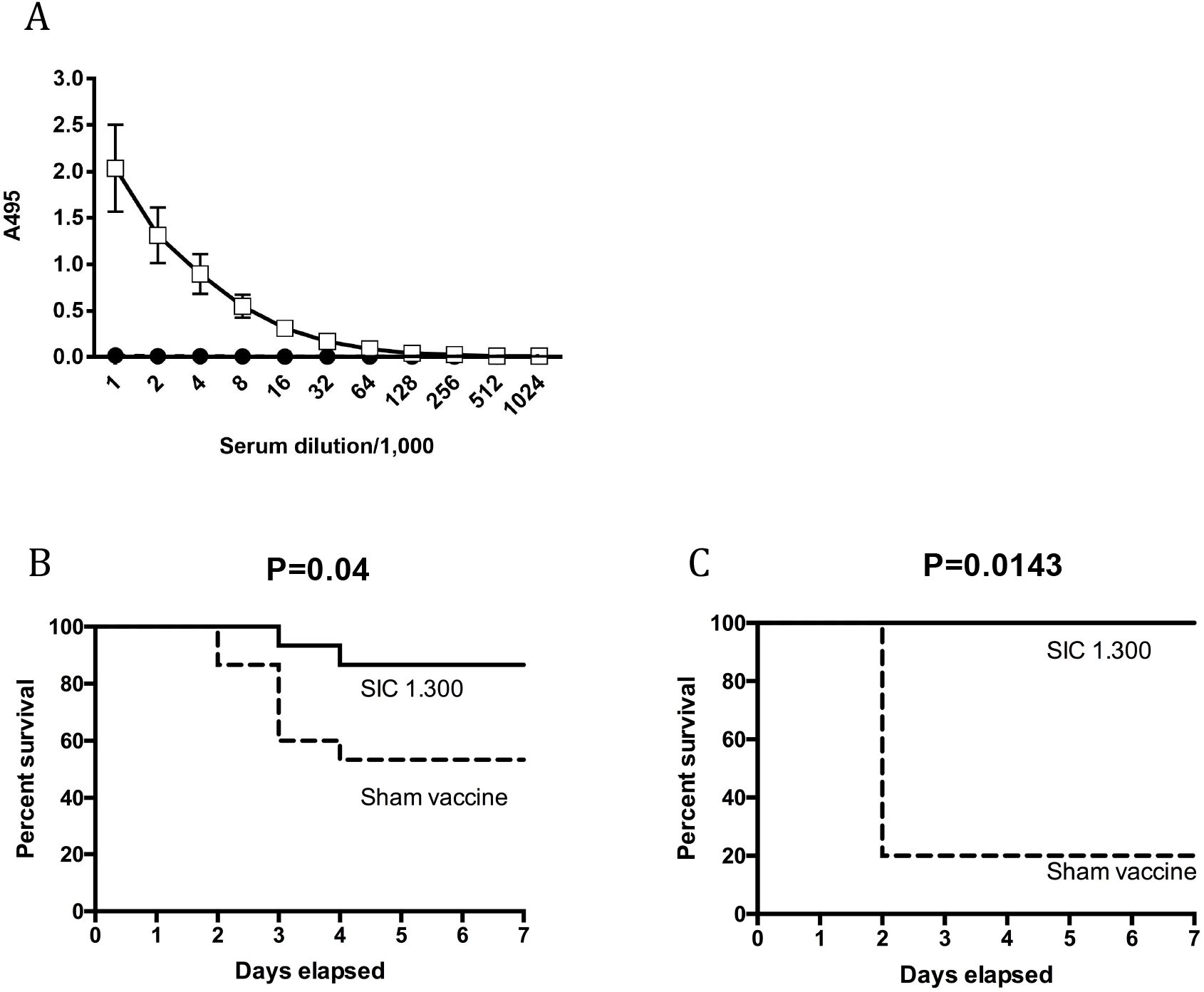
SIC 1.300 vaccination is immunogenic and induces protective response against lower respiratory tract infection. A) Serum was obtained from mice on day 39 post-immunization with rSIC1.300 (open squares) or PBS (closed circles) and SIC1.300-specific IgG was measured by ELISA. Data were obtained from 10 mice per group, over three vaccination experiments. (B-C) FvB/n mice immunized with SIC 1.300 (solid line) or sham vaccinated (dashed line) were infected intranasally with B) 2×10^7^ CFU (n=15, pooled data from two independent experiments) or (C) 2×10^8^ CFU (n=5) of the *emm1 S. pyogenes* isolate H584 and culled when experimental endpoints were reached. Survival was compared using the log-rank test.

### Immunisation with SIC protects against systemic bacterial dissemination

To determine the mechanism of protection, in a separate experiment SIC-immunised mice were challenged intranasally and bacterial counts at the site of infection and in distant tissues were quantified 48 hours after infection (Figure 3). Bacterial counts recovered from the nose were similar between both sets of animals, indicating that there was no differences in dose or local bacterial replication between the two groups (Figure 3A). Compared to mice that received sham vaccination, mice immunised with SIC1.300 had significantly reduced bacterial counts in the spleen (Figure 3B) and liver (Figure 3C). Although 3/10 SIC-immunised mice were bacteraemic, compared to 7/10 sham immunised mice, there was no significant difference in bacterial counts in the bloodstream (Figure 3D).

**Figure 3.**
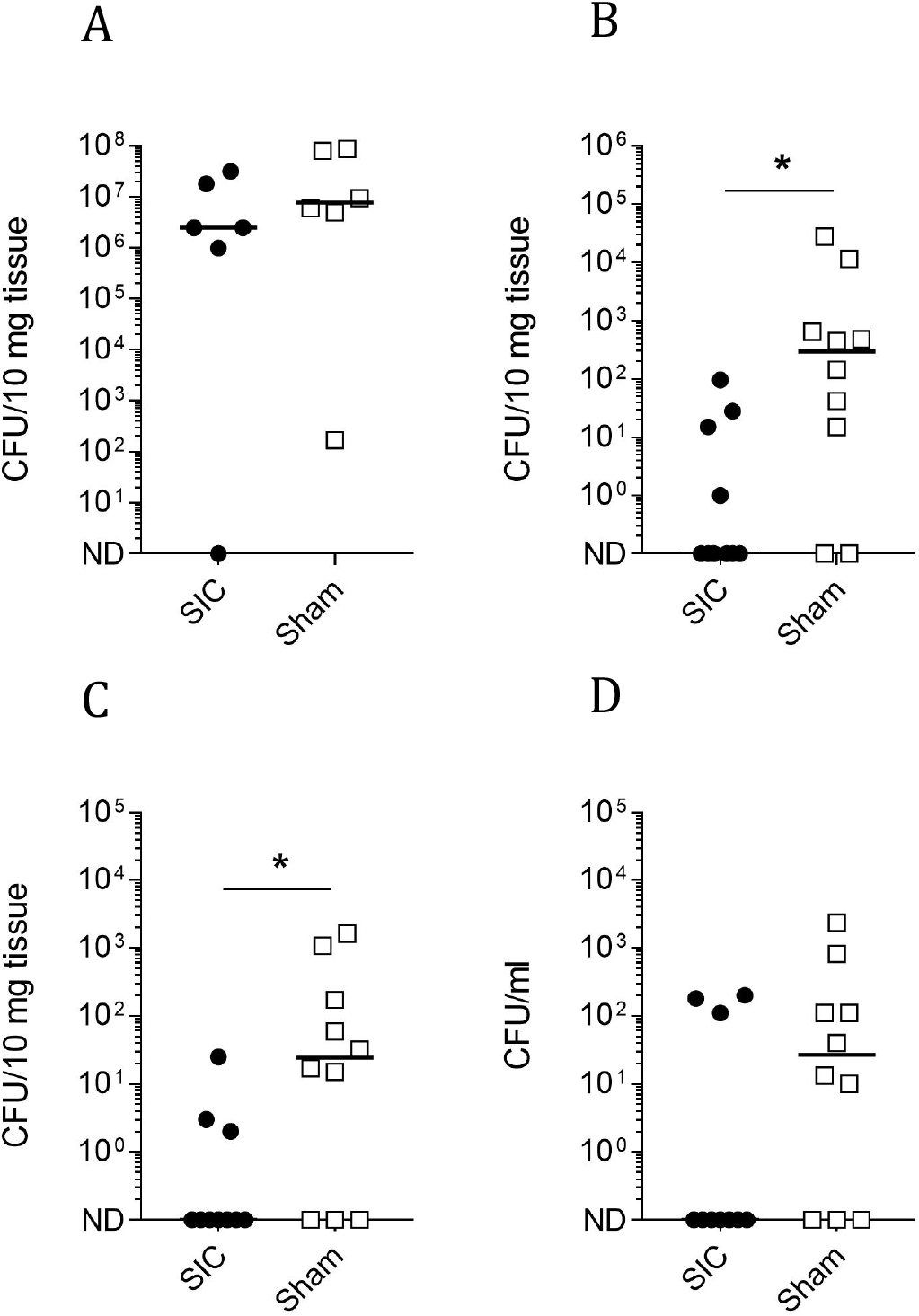
SIC 1:300 vaccination prevents *S. pyogenes* dissemination from an respiratory tract focus of infection. FvB/n mice immunised with SIC 1.300 (n=10) or sham vaccine (n=10) were infected i.n. with 5×10^7^ CFU of the *emm1 S. pyogenes* isolate H584. Mice were culled 48 h post-infection and bacterial loads within the nose (A), spleen (B), liver (C) and blood (D) were enumerated. For the nasal tissue, bacterial enumeration was only performed on 6 mice per group. Solid lines indicate the median CFU recovered from each organ. *p < 0.05 one-tailed Mann-Whitney U. ND: Not Detected.

A separate group of SIC-immunised and sham-immunised mice were infected by the intramuscular route and bacterial counts were assessed 24 hours after infection. Again, while no differences in bacterial load were observed at the site of inoculation, in the infected muscle, (Figure 4A) a significant difference in the bacterial counts in the spleen (Figure 4B) and liver (Figure 4C) was measured. Very limited bacterial dissemination to the blood was observed (Figure 4D).

**Figure 4.**
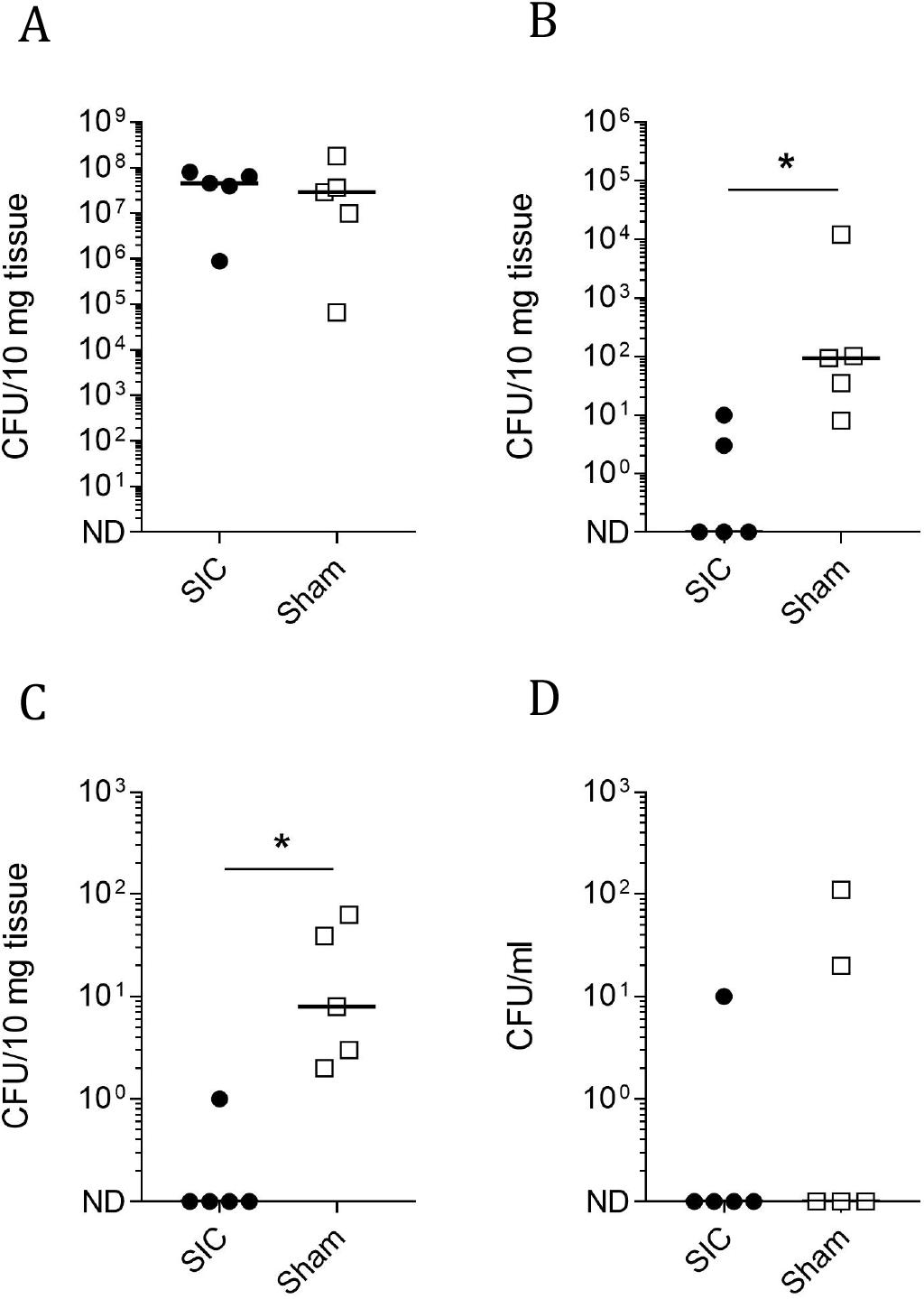
SIC 1:300 vaccination prevents *S. pyogenes* dissemination from an intramuscular focus of infection. FvB/n mice immunised with SIC 1.300 (n=5) or sham vaccine (n=5) were infected I.M. with 5×10^7^ CFU of the *emm1 S. pyogenes* isolate H584. Mice were culled 24 h post-infection and bacterial loads within the thigh muscle (A), spleen (B), liver (C) and blood (D) were enumerated. Solid lines indicate the median CFU recovered from each organ. *p < 0.05 one-tailed Mann-Whitney U. ND: Not Detected.

### Vaccine-induced rabbit anti-SIC antibodies, but not naturally occurring human antibodies enhance clearance of *emm1 S. pyogenes ex vivo*

To assess the protective role of vaccine-induced immunity against SIC *ex vivo*, polyclonal rabbit anti-SIC serum was generated using recombinant SIC 1:300. Cross reactivity with other SIC variants and native SIC from *emm1* GAS culture supernatant was confirmed by ELISA and western blotting (Supplementary Figure S1). To determine if vaccine induced rabbit antibodies and/or naturally occurring human antibodies to SIC could enhance clearance of *emm1 S. pyogenes* in the *ex vivo* whole blood assay, rabbit and human anti-SIC antibodies were affinity purified from rabbit polyclonal serum and commercially-available pooled human immunoglobulin (ivIg) respectively. ELISA and western blot analysis confirmed that anti-SIC antibody from both humans and rabbits, that was affinity purified using SIC1.300, was able to detect full length recombinant (r) SIC of three different variants and also native SIC from *emm1 S. pyogenes* culture supernatant, suggesting that immunogenicity was not restricted to a single SIC variant (Supplementary Figure S2).

Purified rabbit and human anti-SIC antibodies were added to human whole blood from healthy individuals and growth of *emm1 S. pyogenes* was assessed over 3h using a modified Lancefield assay. Rabbit anti-SIC antibody reduced growth of the *emm1* isolates H584 and AP1 compared to rabbit IgG isotype control antibody (Figure 5A). In contrast, human anti-SIC IgG did not inhibit growth of the *emm1* isolate H584 in whole human blood, compared to a control antibody (Figure 5B). We further evaluated naturally occurring SIC antibodies using a panel of human sera previously determined to have high anti-SIC titres ^16^. There was no correlation between anti-SIC titre and bacterial growth inhibition when heat-inactivated human serum from antenatal donors ^16^ was co-incubated with *emm1 S. pyogenes* growing in human whole blood (Supplementary Figure S3), further indicating that natural anti-SIC antibodies in human serum do not promote opsonophagocytic killing of *S. pyogenes*. Together with the *in vivo* data from mice, the findings indicated that immunisation of mice or rabbits with SIC results in antibodies that have the potential to protect against *emm1 S. pyogenes* infection. In contrast, naturally occurring human anti-SIC antibodies lacked the protective activity that was observed for vaccine-induced antibody.

**Figure 5.**
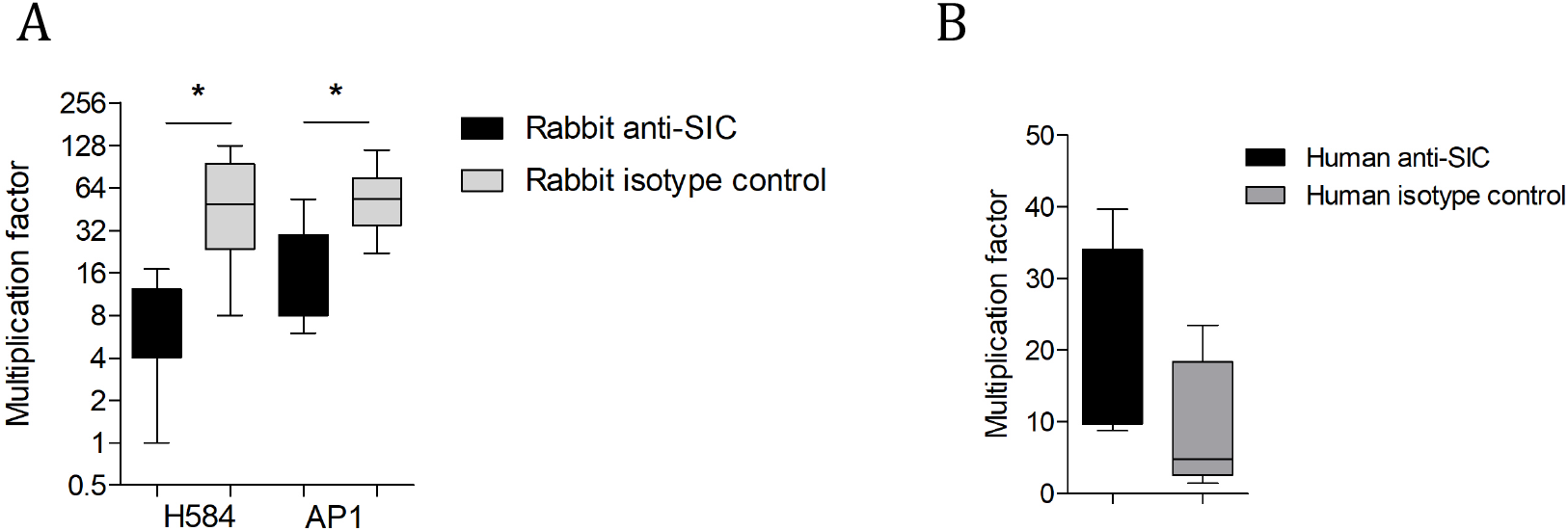
Rabbit but not human anti-SIC antibodies are protective in *ex-vivo* whole blood assay. A) The *emm1 S. pyogenes* isolates H584 and AP1 were grown in human whole blood with the addition of affinity purified rabbit anti-SIC 1.300 polyclonal antibodies (black bars) or rabbit IgG isotype control antibodies (grey bars). Bacterial growth (multiplication factor) was determined after rotation at 37°C for 3 hours. Median and range shown for two separate experiments with three different donors. *p < 0.05 Wilcoxon matched-pairs signed rank test. B) The *emm1 S. pyogenes* isolate H584 was grown in human whole blood with the addition of purified human anti-SIC antibodies (black bars) or human isotype control antibodies (grey bars). Bacterial growth (multiplication factor) was determined after rotation at 37°C for 3 hours. Mean and standard deviation shown for experiment repeated in triplicate from one donor.

### Human, rabbit and mouse anti-SIC antibodies detect different fragments of SIC

To determine the basis for the apparent difference between natural human anti-SIC antibodies, and vaccine-induced rabbit or mouse anti-SIC antibodies, the regions of SIC recognised by each of the anti-SIC antibodies were studied using polypeptide fragments of SIC (fragments 1, 2 and 3), which are based on SIC from the *emm1 S. pyogenes* strain AP1 (Supplementary Figure S4). Whilst purified rabbit anti-SIC antibodies were able to recognise all three SIC fragments by an ELISA-based assay, purified natural human anti-SIC was able to detect only fragment 3, with limited detection of fragment 1 or fragment 2 (Figure 6A). Serum from mice that had been immunised with full length rSIC1.300 for the earlier infection challenge experiments, also detected all three SIC fragments similar to findings in the rSIC1.300-immunised rabbit serum (Figure 6B). These findings were confirmed by Western blot; rabbit and mouse anti SIC detected all three fragments while human anti-SIC detected only fragment 3 (Figure 6C). Furthermore, when the anti-SIC response was quantified from individual donor human antenatal sera, the predominant response was against SIC fragment 3 (Figure 6D).

**Figure. 6.**
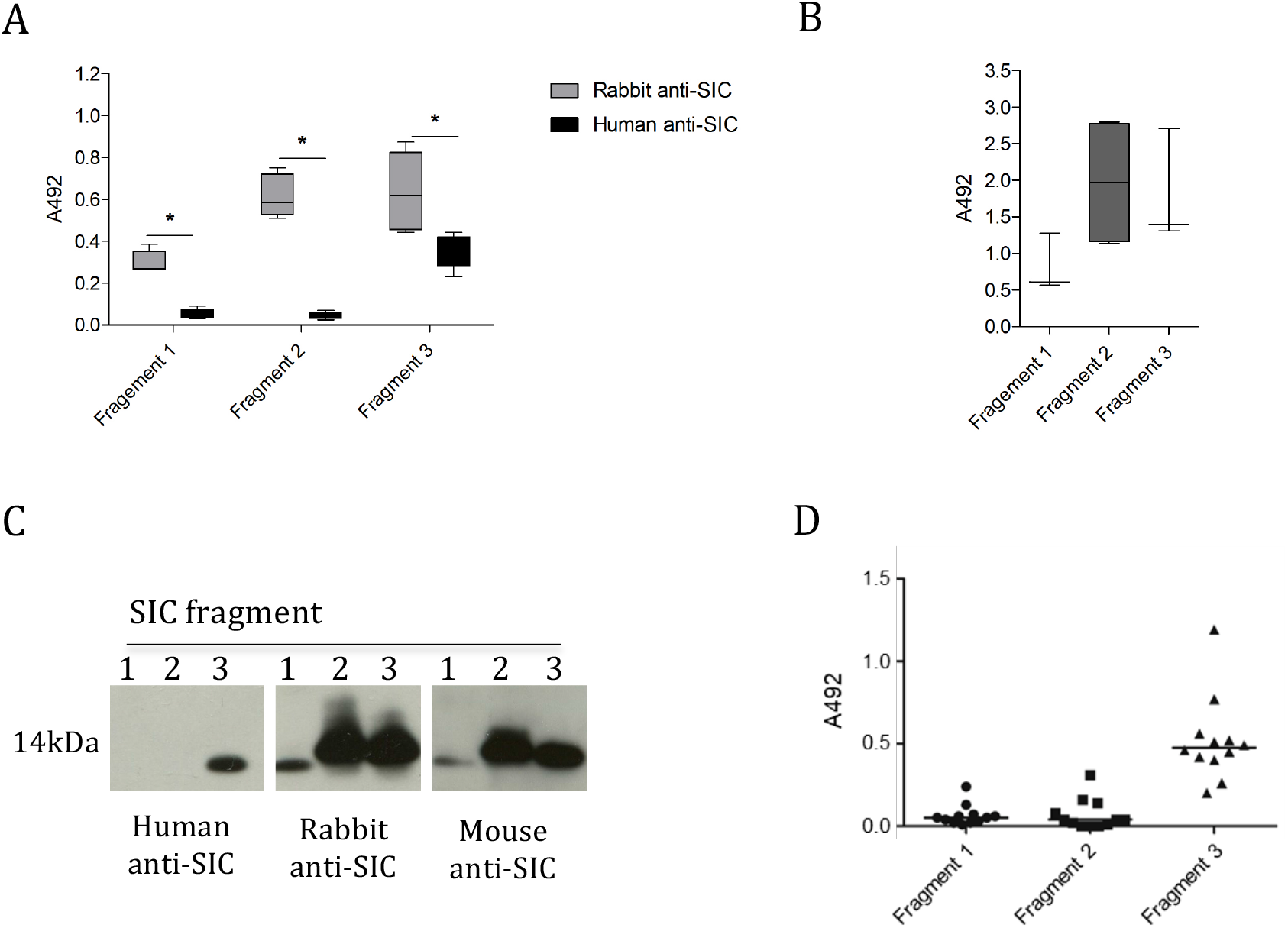
Rabbit and mouse but not human anti-SIC antibodies recognise all three SIC fragments. A-B) Immobilised recombinant SIC fragments 1, 2 and 3 were incubated with A) 0.1mg / ml of affinity purified human anti-SIC or rabbit anti-SIC1.300 antibodies, or B) a 1:100 dilution of pooled serum from mice immunised with full length SIC1.300. Mean and standard deviation shown of ELISAs repeated at least twice. C) Equal quantities of recombinant SIC fragments 1, 2 and 3 were visualized by western blotting using 0.1mg/ml of human anti-SIC or rabbit anti-SIC antibodies, or a 1:250 dilution pooled serum from mice immunised with full length SIC1.300. (D) Immobilised recombinant SIC fragments 1, 2 and 3 were incubated with a 1:100 dilution of heat inactivated sera from individual antenatal donors in which anti-SIC 1.300 titers had been determined previously. Data points represent mean A492 readings from two independent experiments and solid lines indicate the median.

### Immunisation with SIC fragments does not protect against invasive disease

To ascertain whether immunity to any single SIC fragment would be sufficient to induce protective immunity, mice were immunised with recombinant SIC fragment 1, fragment 2, fragment 3 or a sham vaccine containing PBS and adjuvant. Fragment 3 immunization was complicated by an unexpected hypersensitivity reaction in 5/8 mice. Following immunisation, strong reactivity was obtained against SIC fragment 1 and fragment 3 respectively with serum from mice immunised with the homologous SIC fragment when analysed by ELISA (Figure 7A) and western blotting (Figure 7B). No antibodies against SIC fragment 2 were detected by either method.

**Figure 7.**
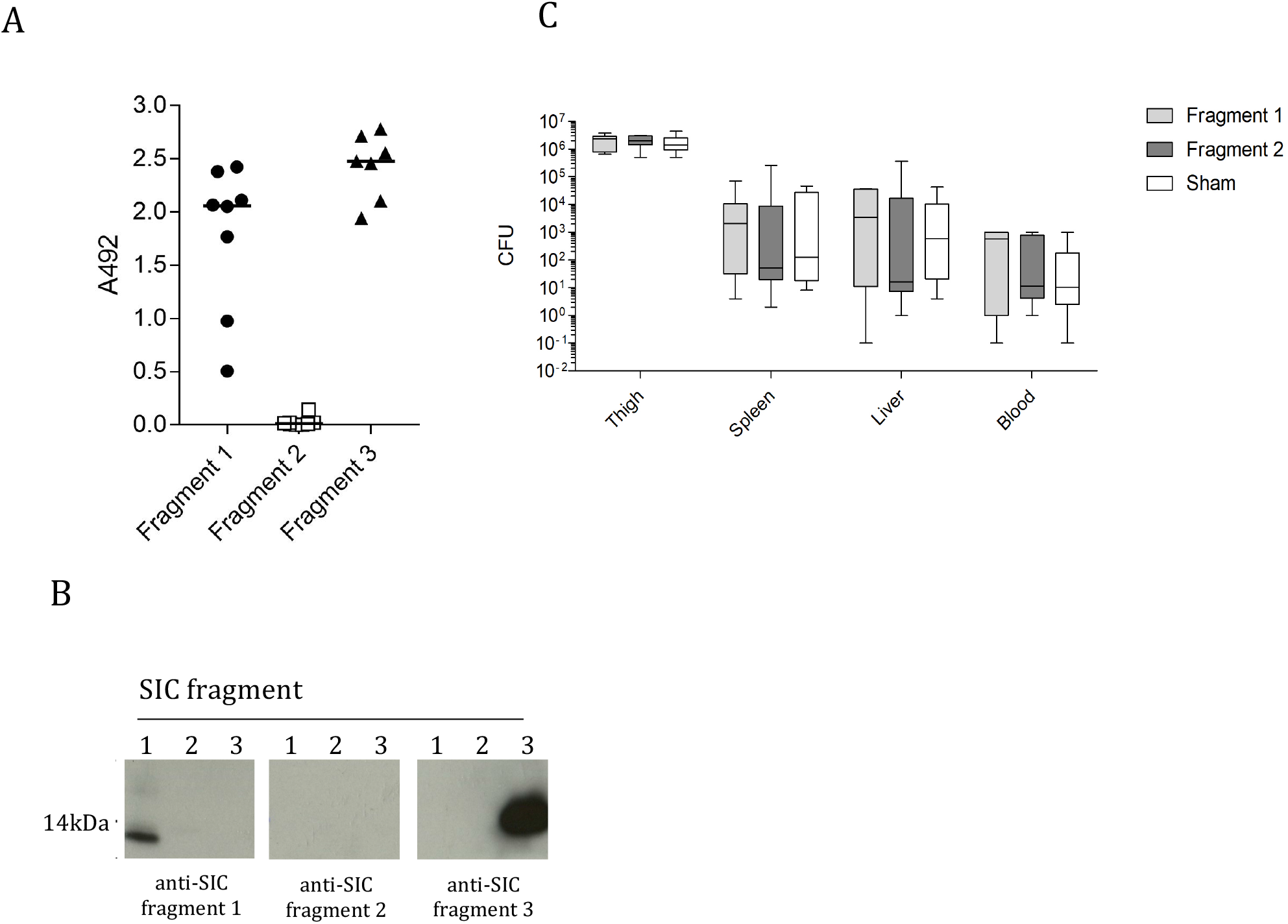
SIC fragments are variably immunogenic and non-protective. A) Sera were obtained from individual mice (n=8) immunized with recombinant SIC fragments 1, 2 or 3 and SIC fragment specific IgG was measured by ELISA. Data points represent mean A492 readings from individual mice obtained in two independent experiments and solid lines indicate the median. B) Equal quantities of recombinant SIC fragments 1, 2 and 3 were transferred to a membrane and incubated in in 1:100 dilution of mouse anti-SIC fragment 1, 2 or 3 antiserum. C) FvB/n mice immunised with SIC fragment 1, fragment 2 or sham vaccine (n=8) were infected intramuscularly with 2 × 10^8^ CFU of the *emm1 S. pyogenes* isolate. Mice were culled 24 h post-infection and bacterial loads within the thigh muscle spleen, liver and blood were enumerated. Data are displayed as CFU/10 mg tissue (thigh, spleen and liver) or CFU/ml (blood) (median and range).

Mice immunised with individual SIC fragments were challenged with 2 × 10^8^ CFU of *emm1 S. pyogenes* AP1 by the intramuscular route and the bacterial counts in the organs and at the site of infection were quantified 24 hours after infection. There was no significant difference in dissemination to the bloodstream, liver, spleen or at the site of infection between any of the groups (Figure 7C). Thus, the protection conferred against *S. pyogenes* by immunisation with full length rSIC could not be recapitulated by any single SIC domain.

## Discussion

Antibodies against SIC are widespread in populations worldwide ^14-16^, although the role that these antibodies play in protection against infection with *S. pyogenes* remains unclear. In this present study, we have demonstrated that SIC is expressed ubiquitously by invasive and non-invasive *emm1* bacterial strains *in vitro* and we have quantified SIC production *in vivo*. Immunisation of animals with full length SIC provided protection against systemic bacterial dissemination. In contrast to vaccine-induced immunity, natural human immunity to SIC is directed against only one domain of SIC, and this is insufficient to confer immunity.

Multiple functions have been attributed to SIC which all appear to aid bacterial evasion of host innate immunity ^6,8,9^. The upregulation of *sic* by invasive *emm1* isolates that had undergone a mutation in the bacterial two-component regulator *covRS*^11^ suggested that SIC may contribute to *S. pyogenes* invasiveness as part of the *covRS* regulon. Whilst previous studies have examined the role of the variation in sequence and size of SIC ^17-21^, we instead hypothesised that variation in SIC expression levels would reflect the invasive phenotype of a strain. SIC expression by single bacterial isolates has been quantified in two separate reports ^8,9^, however, to our knowledge this is the first report to examine SIC expression in a wider collection of invasive and non-invasive *emm1* GAS clinical isolates. Although SIC expression levels varied widely, levels were not overall significantly higher amongst invasive isolates. Of note, invasive isolates represented almost two thirds of the strains investigated and it is likely that only some have mutations in covRS. Among a subset of isolates for which covRS sequencing was undertaken we did however observe significantly heightened expression of SIC. We have, for the first time, also demonstrated that SIC is detectable in infected murine thigh tissue, which was the site of infection, in infected animals. We attempted to detect SIC in muscle from a patient with necrotising myositis caused by *emm1 S. pyogenes*, however samples were obtained after treatment with intravenous immunoglobulin, and showed too much cross reactivity with reagents used, precluding detection of SIC (data not shown). Recently SIC was one of just 15 *S. pyogenes* proteins detected in pleural fluid from a child with empyema caused by *emm1 S. pyogenes*, underlining the potential for virulence factors such as SIC to be upregulated during *in vivo* infection.

Antibodies against SIC are widespread and persistent amongst human populations in geographically distinct regions ^14-16^. Intriguingly, anti-SIC antibodies are identified more frequently than antibodies to M1 protein (14). We set out to assess whether immunisation with one SIC variant (SIC1.300) would be protective *in vivo* in two mouse models of infection. rSIC immunised mice had increased survival compared to sham-immunised mice following respiratory tract infection. Whilst there was no difference in bacterial counts in the nasal tissue and lungs, immunity to SIC appeared to impact bacterial dissemination beyond the site of infection; the observation that rSIC immunisation also reduced bacterial dissemination from intramuscular infection provided further evidence to support this.

The ability of antibodies raised against full length SIC to enhance clearance of *S. pyogenes* in whole blood and in the mouse models is likely to reflect inhibition of SIC function rather than any opsonic activity, since we found little convincing evidence of surface-localised SIC. The data are consistent with previous work demonstrating the role of SIC *in vivo*: SIC enhanced virulence in both intraperitoneal ^12^ and subcutaneous mouse models of GAS infection ^9,22^. By immunising mice with SIC and demonstrating reduced bacterial dissemination following infectious challenge, our data suggest an important role for SIC in *emm1 S. pyogenes* pathogenesis and, based on observations in the whole blood model, a potential role in resistance to opsonophagocytic killing.

Whilst human anti-SIC antibodies are abundant in populations, our data suggests that these antibodies are not protective. One possible explanation for this is that following immunisation with full length SIC, protective antibodies are raised to epitopes throughout the mature SIC protein, however, following natural infection, antibodies are only generated against epitopes within fragment 3 of SIC. A previous study using sera from 29 individuals, identified ten linear epitopes in SIC1.01 that were identified by ≥ 50% of the human anti-SIC sera. These epitopes were at sites in SIC in which polymorphisms commonly occur, and were evenly distributed within the equivalent of SIC fragments 1, 2 and 3 ^14^. When this was further analysed by phage display using two sera to detect natively folded SIC peptides, 7/8 peptides recognised by one serum and 11/12 peptides recognised by the other serum spanned regions present in the equivalent of SIC fragment 3 (from residue 168 onwards). Our data using ELISA and western blot confirm that epitopes within fragment 3 are the most readily recognised by human anti-SIC that had been purified from ivIg pooled from over 1,000 donors 23

The reasons that human anti-SIC antibody responses are directed to a single domain only are unclear. One possibility is that anti-SIC responses represent cross-reactive responses to another, unrelated antigen, that has structural similarity to fragment 3. Notably, SIC readily undergoes proteolysis by human proteases such as human neutrophil elastase and bacterial proteases such as

SpeB ^14,24^. The digestion of SIC by unknown bacterial or host factors within human saliva ^25^ provides an alternative mechanism by which SIC fragments, rather than full length SIC, might be presented to the immune system.

The lack of protection following immunisation with separate SIC fragments was surprising, especially considering that several immune inhibitory functions of SIC have been localised to the SRR and LRR contained within SIC fragments 1 and 2 respectively ^5^. Indeed, immunisation with SIC fragment 2 elicited minimal antibody response, despite epitopes in this fragment being detected with antibodies raised against full length SIC, and also human anti-SIC serum in a previous study ^14^, suggesting that this fragment may form part of a discontinuous epitope. Data generated using various biophysical methods indicate that SIC contains low levels of regular secondary and tertiary structures (unpublished data); this could mean that discontinous epitopes form long range contacts resulting in either stable or dynamic tertiary structure assemblies. Resolution of the complete folded structure of SIC would provide a better understanding of the nature of these epitopes. Interestingly, few immune inhibitory functions of SIC have been localised to the PRR of SIC contained within fragment 3. Thus, an alternative explanation to the varying immunogenicity of SIC domains is that, in humans, SIC fragments 1 and 2 bind to host ligands concealing these regions from the host, whilst epitopes in SIC fragment 3 are abundantly available. Whilst SIC binds to both human LL-37 and mouse cathelicidin ^12^, previous studies assessing other ligands of SIC have used only human proteins ^5,8,9,22,24^. It remains unclear whether SIC binds to other mouse proteins and hence the lack of immunogenicity of fragment 2 compared to fragment 1 may be due to differential binding to mouse proteins.

At first glance, a vaccine based on an antigen found in only one serotype, does not seem an attractive proposition. However, in light of difficulties in developing an effective GAS vaccine over the last 70 years, and the dominance of the *emm1* lineage, alternative approaches must be considered. The successful introduction of multi-component vaccines for other bacterial infections indicates that the inclusion of multiple novel antigens in a vaccine may be important. Additionally, antibodies directed against single virulence factors are being developed as adjunctive therapy for a number of serious diseases caused by other bacteria. Our data suggest that the inclusion of SIC in a multi-component vaccine has potential to reduce the burden of disease caused by highly invasive *emm1 S. pyogenes* and goes someway to explain the anomaly of widespread anti-SIC immunoreactivity in an otherwise susceptible population.

## Materials and Methods

### Bacterial strains

*S. pyogenes emm1* strain H584 was isolated from a case of lethal postpartum sepsis ^16^. An additional 100 *emm1* clinical *S. pyogenes* isolates from the 1930s through to 2013 were referred to Imperial College from diagnostic laboratories, linked to available clinical data, and anonymised as approved by the local research ethics committee (06/Q0406/20). 68 isolates were from invasive disease and 32 were from non-invasive disease. Specific disease etiologies were not known for 13 isolates. *S. pyogenes* isolates were identified as *emm1* via sequencing of the *emm* gene, which encodes the M protein (https://www2a.cdc.gov/ncidod/biotech/strepblast.asp). Invasive disease-associated isolates were defined as *S. pyogenes* isolated from a normally sterile site or a non-sterile site with clinical diagnosis of necrotising fasciitis or septic shock. Non-invasive disease-associated isolates were defined as *S. pyogenes* isolated from non-sterile sites with no clinical signs indicating severe/invasive disease. *S. pyogenes emm1* strain AP1 and SIC-, a mutant derivative not expressing SIC, were used in experiments where a SIC negative *emm1* isolate was required and have been described previously ^9^.

All streptococcal strains were cultured on Columbia horse blood agar (CBA) (E&O Laboratories Ltd, Bonnybridge, Scotland) or in Todd-Hewitt broth (THB) (Oxoid, Basingstoke, UK) at 37 °C, 5% CO2 without shaking. *Escherichia coli* were cultured in Luria-Bertani broth (Oxoid) with 100 μg/ml ampicillin (Sigma-Aldrich, Dorset, UK).

### Recombinant SIC proteins

Recombinant full length SIC proteins were purified using affinity chromatography with a His-bind column as per manufacturer’s instructions (Novagen, Merck, Darmstadt, Germany). *sic*1.300, from *emm1* strain H584, *sic*1.02 and *sic*1.301, used in initial validation experiments were expressed in *E. coli* using the pET19b expression vector (Novagen, Nottingham, UK), as previously described ^16^. Recombinant SIC fragment 1 (amino acids 1-69), fragment 2 (amino acids 70-167) and fragment 3 (amino acids 168-278) were based on the originally published *sic* sequence of the *emm1* strain AP1 ^4^ and were expressed in *E. coli*, purified by nickel affinity chromatography, processed with TEV protease to remove the His-tag, purified by reversed phase chromatography and lyophilized.

### Murine immunisation and infection challenge

Female six to eight-week-old FVB/n mice (Charles River, Margate, UK) were immunised intramuscularly with 30 μg of recombinant SIC protein or recombinant SIC fragments 1, 2 and 3 emulsified 1:1 in Freund’s incomplete adjuvant (Sigma-Aldrich) on days 0, 21 and 35; sera were collected by tail bleed on day 39-41. A control group of mice were immunised with phosphate buffered saline (PBS) and adjuvant (sham vaccination). On day 42-45, mice were infected with *emm1 S. pyogenes* strain H584 after full length SIC immunisation, or *S. pyogenes* strain AP1 following SIC fragment immunisation, (as SIC fragments were derived from SIC AP1). For respiratory tract infection, two invasive disease isolates were used. Mice were briefly anaesthetised with isoflurane and 2 × 10^7^

CFU or 2 × 10^8^ CFU of *S. pyogenes* were administered by inhalation (5 μl of bacterial suspension per nostril). Mice were monitored for seven days and any mice reaching defined humane endpoints were euthanized. For intramuscular infection, mice received 5 × 10^7^ CFU of GAS directly into thigh muscle. In some experiments mice were euthanized (after 48 hours for intranasal infection and 24 hours for intramuscular infection) and blood and tissue (homogenised in PBS) were taken for culture.

### Purification of antibodies from serum or human intravenous immunoglobulin

Rabbit anti-SIC polyclonal serum was raised against recombinant SIC1.300. To purify anti-SIC antibodies from rabbit anti-SIC serum or human pooled intravenous immunoglobulin ((ivIg), Privigen immune globulin, CSL Behring, PA, USA)) recombinant SIC1.300 (1 mg/ml) in coupling buffer (0.1M sodium bicarbonate, 0.5M sodium chloride, pH8.3) was applied to a chromatography column containing CnBr activated agarose (Sigma-Aldrich) at room temperature for 2 hours. Following washing with coupling buffer, 0.2M glycine was applied at room temperature for 2 hours. The SIC-CnBr resin was then washed extensively by alternating between coupling buffer and 0.1M acetate buffer containing 0.5M sodium chloride, pH4. The resin was equlibrated in Tris buffered saline 1 [(TBS1), 50mM Tris, 150mM NaCl, pH 7.5)], then either serum from a rabbit immunised with SIC1.300 or human pooled ivIg was applied to the SIC-CnBr resin. The resin was then washed extensively with TBS1 and Tris buffered saline 2 [(TBS2), 20mM Tris, 2M NaCl, pH7.5). Purified antibodies against SIC were eluted in fractions from the resin using 0.2M glycine-HCl, pH2.2 and neutralised with 1M Tris-HCl, pH 8.0. Fractions containing purified anti-SIC antibodies were pooled and dialysed into PBS overnight at 4°C. Antibody concentrations were determined using the Coomassie-Bradford assay. From 12 ml of rabbit anti-SIC serum, approximately 2 ml of purified rabbit anti-SIC antibody (0.5mg/ml) was obtained, and from 20 ml of human pooled ivIg, approximately 0.5 ml of purified human anti-SIC antibody (0.1mg/ml) was obtained.

### Human whole blood phagocytosis assay

Lancefield whole blood assays were performed to assess the protective effect of rabbit and human anti-SIC antibodies *ex vivo.* Approximately 50 CFU of *emm1 S. pyogenes* strain H584 or AP1 were inoculated into heparinized human whole blood obtained from healthy donors as described ^26^. Mixtures of whole blood, bacteria and anti-SIC were incubated for 3 hours at 37°C with end-over-end rotation. Bacterial survival was quantified as the multiplication factor of number of surviving colonies relative to the starting inoculum and tested in triplicate. To assess the effect of rabbit anti-SIC, purified rabbit anti-SIC antibodies were added to the whole blood. Rabbit IgG isotype control antibody (ab176094, Abcam, Cambridge, UK) was used as a control. For separate studies with H584 using human anti-SIC, either serum from 79 healthy antenatal patients or purified human anti-SIC from human pooled ivIg were added to the whole blood. Antenatal sera were previously used and tested for anti-SIC1.300 titres ^16^. A human IgG isotype control (Novus biological, CO, USA) was used as a control.

### ELISA-based assay

To assess the antibody response of immunised mice, 96-well polystyrene plates (Nunc, ThermoScientific, MA, USA) were coated with 100 ng of full length SIC1.300 or SIC fragments 1, 2 or 3 overnight at 4°C, washed, blocked for one hour with 3% normal goat serum (Sigma-Aldrich) diluted in PBS-0.1% Tween20 (PBST). Test sera were then added at a range of dilutions. Binding was detected using HRP-conjugated goat anti-mouse IgG (Abcam) and incubated for one hour at room temperature. The substrate (ONPG, Sigma-Aldrich) was added to wells, the reaction was stopped with 3N HCl, and the OD A492 read with a μQuant spectrophotometer (Biotek, VT, USA). To compare cross detection of recombinant SIC variants or SIC fragments by rabbit polyclonal anti-SIC1.300 serum or purified SIC antibody, plates were coated with SIC1.02, SIC1.301 and SIC1.300 or SIC fragments 1,2 and 3 and binding was detected using 1 in 25,000 dilution of HRP-conjugated goat anti-rabbit IgG (Life Technologies, Paisley, UK). To determine detection of SIC fragments by human purified anti-SIC antibody or antenatal serum, plates were coated with SIC1.300 or SIC fragments 1, 2 and 3, and binding detected using 1 in 25,000 dilution of HRP-conjugated goat anti-human IgG (Sigma-Aldrich).

### SDS-PAGE and Western blot

Protein samples were separated by sodium dodecyl sulphate polyacrylamide gel electrophoresis (SDS-PAGE) using NUPAGE 10% Bis-Tris Gels (Life Technologies), run in NUPAGE MES Running Buffer (Life Technologies). Following SDS-PAGE, proteins were transferred onto PVDF membranes (Amersham Hybond-LFP membranes [GE Healthcare, Buckinghamshire, UK]), or nitrocellulose membranes (Amersham Protran, GE Healthcare) using cold tris-glycine transfer buffer (0.025M Tris, 0.2M Glycine). For quantification of SIC and cross detection of SIC variants, following blocking in PBST with 5% skimmed milk powder, SIC was detected by probing with 1 in 10,000 dilution of rabbit polyclonal anti-SIC1.300 for two hours at room temperature. Proteins were visualized by incubation in 1 in 50,000 dilution of HRP-conjugated goat anti-rabbit IgG (Life Technologies) for one hour followed by Amersham ECL Prime Western Blotting Detection Reagent (GE Healthcare) exposed to Amersham Hyperfilm ECL (GE Healthcare). For cross detection of SIC variants using purified human anti-SIC, membranes were probed with 1 in 10,000 dilution of human anti-SIC followed by 1 in 50,000 dilution of HRP-conjugated goat anti-human IgG (Sigma-Aldrich). To directly compare detection of SIC fragments by purified human and rabbit anti-SIC antibodies, purified antibodies were first diluted to the same concentration (0.1mg/ml in PBS), then used at 1 in 1,000 dilution to probe membranes, followed by secondary antibodies as outlined. Finally, for detection of SIC fragments by mouse anti-SIC fragment 1, 2 and 3, serum from groups of mice immunised with each fragment was pooled and used to probe membranes at 1 in 100 dilution, followed by 1 in 50,000 dilution of HRP-conjugated goat anti-mouse IgG (Abcam). Full, unmodified western blots for Figures 1C, 6C and 7B are displayed in separate Supplementary materials.

### Quantification of SIC expression in culture supernatant and *in vivo*

GAS strains were grown overnight in THB at 37°C with 5% CO2. Cultures were then centrifuged at 2,500 × g for 10 min at 4°C and proteins from the supernatant were precipitated using 10% tricholoroacetic acid (TCA) with acetone (Sigma-Aldrich). 1,700 μl of TCA-acetone was added to 300 μl of culture supernatant and incubated at −20°C for one hour. Samples were washed twice with ice-cold acetone and allowed to dry before being re-suspended in 30 μl of sample treatment buffer [Lithium Dodecyl Sulphate Sample Buffer (Life Technologies); 100mM DL-Dithiothreitol, (Sigma-Aldrich)], heated at 70°C for 10 minutes, and used for Western blot analysis.

For *in vivo* determination of SIC, female six to eight-week old FVB/n mice (Charles River, Margate, UK) were infected intramuscularly with either 5 × 10^7^ CFU of *emm1 S. pyogenes* strains AP1 or SIC- and culled 3 hours after infection. Infected thigh muscle was homogenised in PBS and centrifuged at 16,000 xg for five minutes. The supernatant was added to sample treatment buffer, heated at 70°C for 10 minutes, and used for Western blot analysis. Serial dilutions of known quantities of SIC1.300 were included on each gel to generate the standard curve, and densitometry was undertaken with Image J software (National Institutes of Health, MD, USA) enabling pixel values to be converted to concentrations.

### Statistical Analyses

Non-parametric tests were used for comparisons and were performed with GraphPad Prism 6.0 software (GraphPad Software, CA, USA). Samples sizes were selected based on pilot studies or previous studies ^27^. No randomisation or blinding was performed. All animal procedures were approved by the local ethical review process and conducted in accordance with the relevant, UK Home Office approved, project license (PPL70/7379).

### Study approval

The analysis of anonymised samples and bacteria from patients with suspected infection was approved by an NHS Research Ethics Committee (REC reference 06/Q0406/20). Human blood cells from normal donors was obtained following informed consent from a sub-collection of the Imperial College Healthcare NHS Trust Tissue Bank. All animal procedures were conducted in accordance with UK Home Office guidance and approval.

## Acknowledgements

This work was support by the Wellcome Trust (LKKT), UK NIHR Biomedical Research Centre funding scheme (SS), the Swedish Research Council, project 7480 (LB, IMF), the Knut and Alice Wallenberg Foundation (LB), the Alfred Österlund Foundation (IMF,LB), Hansa Biopharma (LB), the Swedish Government Funds for Clinical Research, ALF (LB).

## Competing Interest Statement

The authors have declared that no conflict of interest exists

## Authors’ contributions

LKKT, MR, LB and SS conceived the study. LKKT and MR analysed the data and wrote the manuscript. LKKT, MR, DT, NR, VN, LEML, CT, IMF and MW performed the experiments. LKKT and MR prepared the Figures. All authors reviewed and approved the final version of the manuscript.

## Supplementary figures

**Supplementary figure S1.**
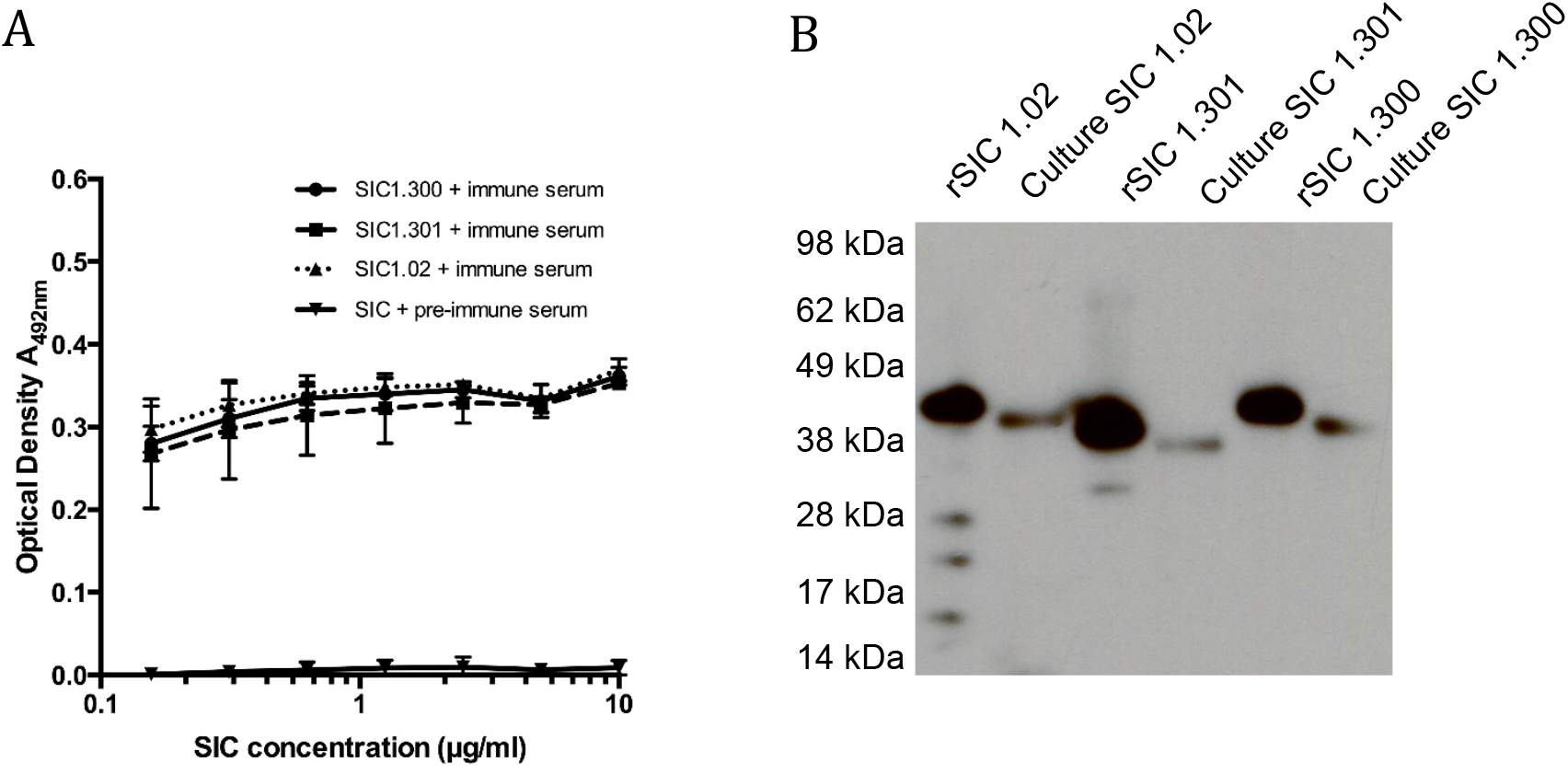
Rabbit anti-SIC1.300 cross detects other SIC variants. (A) Increasing concentrations of different recombinant (r)SIC variants (rSIC1.02 dotted line, triangles; rSIC1.300 solid line, circles; and rSIC1.301 dashed line, squares) were bound to ELISA wells and incubated with 1:10,000 dilution rabbit anti-SIC1.300 serum. Each recombinant SIC variant bound to ELISA wells was also separately incubated in pre-immune rabbit serum (triangles). Mean and standard deviation shown of experimental triplicates. (B) Recombinant SIC variants rSIC1.02, 1.300,1.301 and proteins from concentrated culture supernatant of *S. pyogenes* isolates naturally expressing the same SIC variants were transferred to a membrane and incubated with 1:10,000 dilution of rabbit anti-SIC1.300. SIC1.301 differs from SIC1.300 by a 29 amino acid deletion in SIC1.301, within the long repeat region. SIC1.02 is the common SIC allele in the UK, and differs from both SIC1.300 and SIC1.301 by a 5 amino acid insertion and an amino acid substitution of glutamine (Q) to lysine (K) in the short repeat region ^16^.

**Supplementary figure S2.**
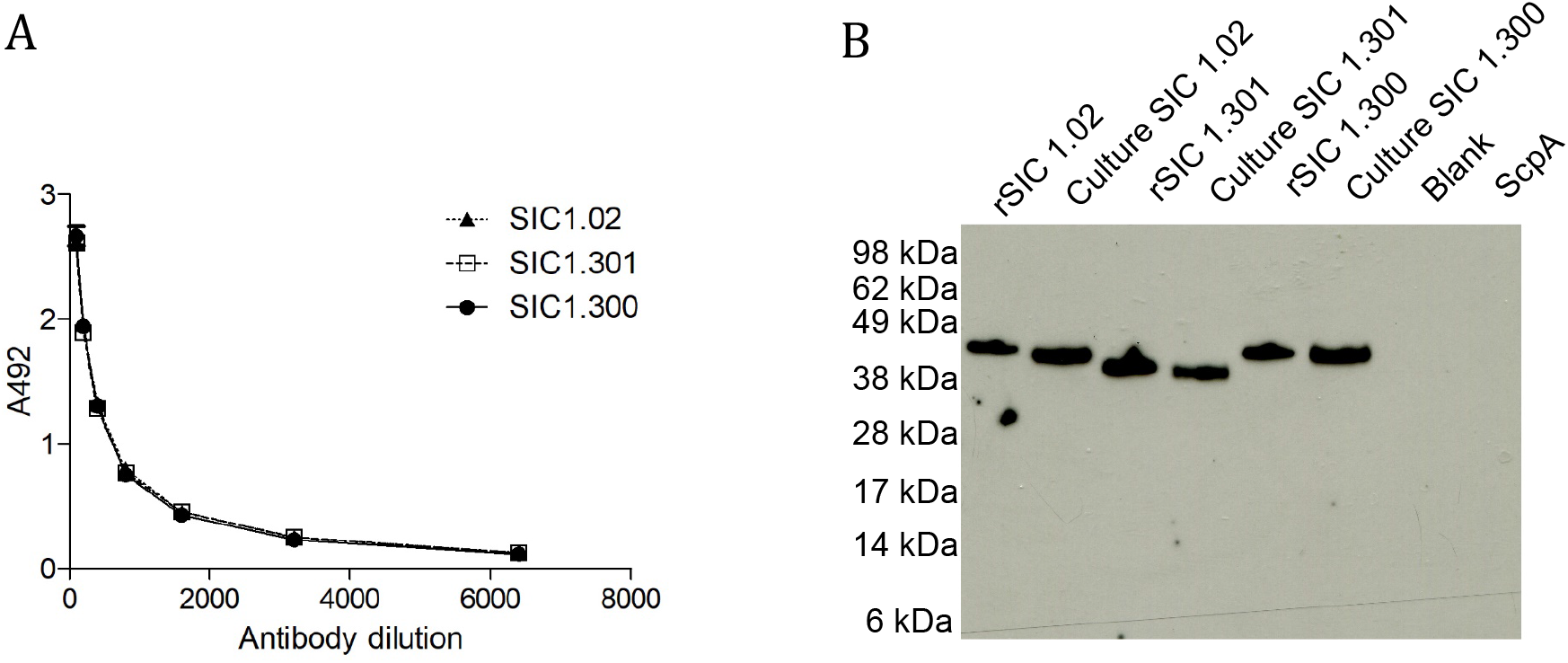
Purified human anti-SIC cross-detects SIC variants. (A) Affinity purified human anti-SIC antibodies were obtained from pooled human intravenous immunoglobulin and SIC-specific IgG against rSIC1.02 (dotted line, triangles), rSIC1.300 (solid line, circles), rSIC1.301 (dashed line, squares) was measured by ELISA. Data show the mean and standard deviation from experimental triplicate. (B) Recombinant SIC variants (25 ng) rSIC1.02, rSIC 1.300, rSIC 1.301, and proteins from concentrated culture supernatant (10 μl) of *S. pyogenes* isolates naturally expressing the same SIC variants were transferred to a membrane and incubated in 1:1,000 dilution of purified human anti-SIC antibodies. Recombinant streptococcal protein ScpA acted as a negative control.

**Supplementary figure S3.**
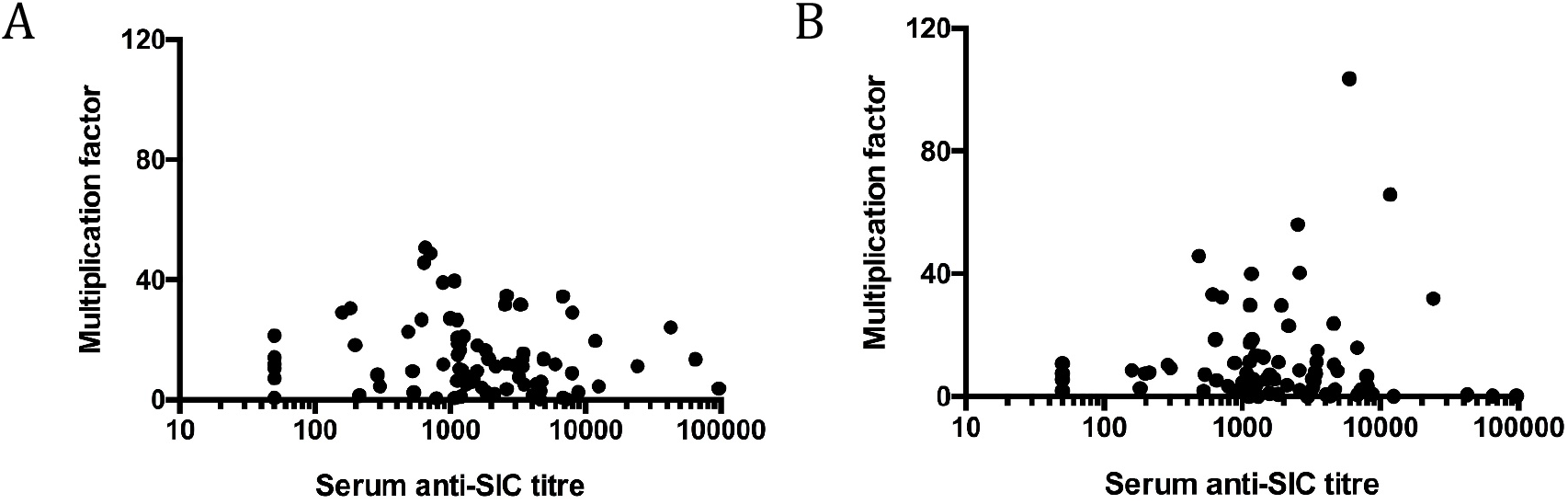
Lack of correlation between human serum anti-SIC levels and bacterial killing. *S. pyogenes emm1* strain H584 was grown in human whole blood from two different donors (A) and (B) co-incubated with individual heat inactivated sera from antenatal donors (n=79) in which anti-SIC 1.300 titres had been previously determined. Bacterial growth (multiplication factor) was analysed after rotation at 37°C for 3 hours and correlated with anti-SIC serum titres. There was no correlation between anti-SIC titres and bacterial growth (Spearman rank coefficient) for donor A (p=0.120) or donor B (p=0.2487).

**Supplementary figure S4.**
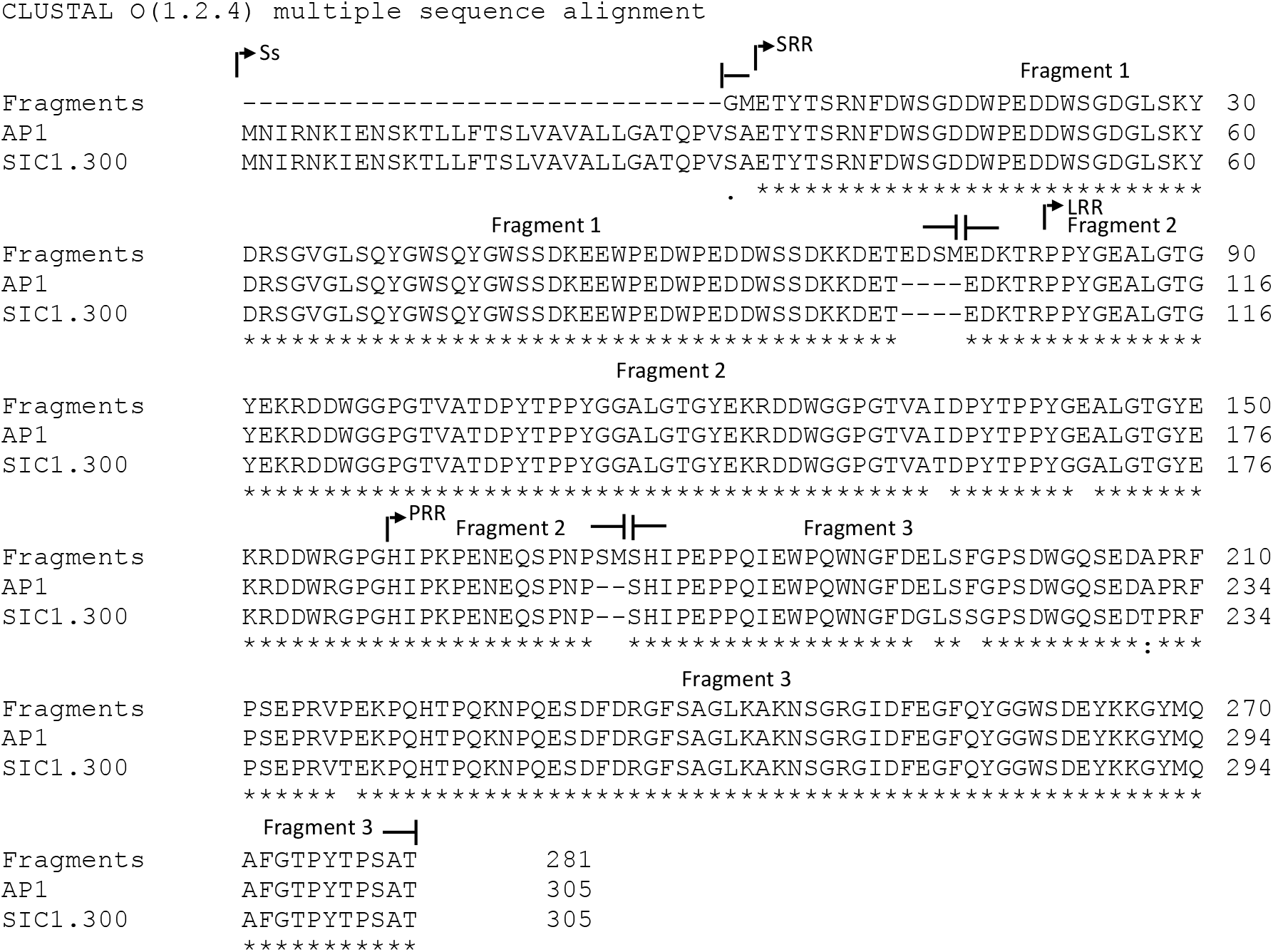
Alignment of amino acid sequences for SIC fragments 1, 2, and 3, SIC 1.300 and SIC from AP1 *emm1* GAS. Recombinant SIC fragment 1 (amino acids 1-69), fragment 2 (amino acids 70167) and fragment 3 (amino acids 168-278) were based on published *sic* sequence of *emm1* strain AP1 ^4^. Arrows mark the start of key regions of SIC: the Signal Sequence (Ss); NH2-terminal short repeat region (SRR); Long repeat region (LRR); Proline rich region (PRR). SIC fragment 1 corresponds to the SRR, fragment 2 corresponds to the LRR and the first 13 amino acids of the PRR, and fragment 3 corresponds to the remainder of the PRR. The start of a SIC fragment is delineated by ⊢ and the end of a fragment is delineated by ⊣. Regions of differences between SIC fragments 1, 2 and 3, SIC1.300 and AP1 SIC are indicated with spaces, dashes (-) indicating absent amino acids, stars (*) indicate identical amino acids between variants and colons (:) indicate equivalent but not the same amino acids. Alignment made using Clustal Omega software and sequences accessed from GenBank.

## Notes

### Competing Interest Statement

The authors have declared no competing interest.

